# Human Cytomegalovirus UL78 is a Nuclear-Localized GPCR Necessary for Efficient Reactivation from Latent Infection in CD34^+^ Hematopoietic Progenitor Cells

**DOI:** 10.1101/2025.06.02.657350

**Authors:** Samuel Medica, Nicole L. Diggins, Michael Denton, Rebekah L. Turner, Lydia J. Pung, Adam T. Mayo, Jennifer Mitchell, Luke Slind, Linh K. Nguyen, Teresa A. Beechwood, Gauthami Sulgey, Craig N. Kreklywich, Daniel Malouli, Patrizia Caposio, Daniel N. Streblow, Meaghan H. Hancock

## Abstract

Human cytomegalovirus (HCMV) is a ubiquitous pathogen that persists throughout the lifetime of the host due in part to the establishment of latency in CD34^+^ hematopoietic progenitor cells (HPCs) and CD14^+^ monocytes. HCMV encodes four putative G protein-coupled receptors (GPCRs): US27, US28, UL33, and UL78. While the roles of most of these receptors have been investigated, a definitive role for UL78 in HCMV infection has yet to be elucidated. Utilizing an *in vitro* CD34^+^ HPC model, we demonstrate that a recombinant virus lacking UL78 protein expression fails to efficiently reactivate from latent infection. Furthermore, we show using a Lumit-based assay that UL78 preferentially couples to the Gα_i_ family of G proteins and that a recombinant HCMV containing mutations in the UL78 G protein-coupling DRL motif also fails to reactivate from latent infection. Together our findings indicate that Gα_i_ coupling is important for UL78 function during reactivation in latently infected CD34^+^ HPCs, however the protein is not required to establish or maintain latency. To better understand the role of UL78, we conducted proximity-dependent labeling analyses in HCMV-UL78-TurboID infected fibroblasts and CD34^+^ HPCs undergoing reactivation from latency. Congruent with our coupling data, we found Gα_i_ was the only heterotrimeric Gα protein in proximity to UL78. Pathway analysis of the UL78 interactome revealed proteins associated with membrane trafficking, signaling, and the nuclear pore complex as enriched in both cell types. In addition, the UL78 interactome contained viral proteins with nuclear localization including viral transcription and DNA replication machinery. Nuclear localization of UL78 was validated using cell fractionation, immunofluorescence microscopy, and proximity-dependent labelling of isolated nuclei. Together, our results provide novel insights into the localization and function of UL78, previously unknown to contribute to reactivation from latent infection.

**Author Summary:** Human cytomegalovirus (HCMV) remains one of the most widespread viral infections globally. Primary HCMV infection is typically asymptomatic and leads to the establishment of latency in myeloid lineage cells, where the virus persists for the host’s lifetime. Reactivation of latent HCMV can cause severe complications, particularly in immunocompromised individuals such as transplant recipients and people living with HIV. Several factors influence the transition from latent to lytic infection, including signal transduction through the viral G protein-coupled receptors: US27, US28, UL33, and UL78. Using an advanced *in vitro* model, we show that recombinant viruses lacking UL78 fail to efficiently reactivate from latent infection. Moreover, we show that UL78 preferentially couples to the Gα_i_ family of G proteins via a conserved DRL motif, and this coupling is required for efficient reactivation. These results were confirmed by proximity-dependent labeling experiments where we identified Gα_i_ and several other proteins involved in trafficking, signaling, transcription, and nuclear localization. Nuclear localization of UL78 was confirmed by cell fractionation, immunofluorescence microscopy, and proximity-dependent labeling in isolated nuclei. Collectively, our results uncover a novel role for UL78 in reactivation from latency and shed new light on its localization and function.

## Introduction

Cytomegaloviruses (CMVs) are species-specific herpesviruses that establish life-long infections in their hosts. Human CMV (HCMV) achieves persistence in part through the ability to establish latent infections in CD34^+^ hematopoietic progenitor cells (HPCs) and CD14^+^ monocytes (1,2). Clinical reactivation of latent virus can occur in situations of immunosuppression, such as during allogenic or solid organ transplantation (3). Significant morbidity and mortality is associated with HCMV reactivation following transplantation, and currently available antiviral treatments targeting the DNA replication machinery have toxic side effects that can exacerbate disease and lead to the emergence of drug-resistant variants (4–6). Targeting the latent reservoir and/or early reactivation events is an alternative approach that requires detailed knowledge of the viral and cellular factors that regulate these processes.

HCMV encodes four G protein-coupled receptors (US27, US28, UL33, and UL78) that are thought to mimic the functions of cellular chemokine receptors (7). While the functions of US27, US28 and UL33 have been interrogated in the context of lytic and latent infection, much less is known about UL78 (8–12). The UL78 family includes HCMV UL78, rat CMV (RCMV) R78, murine CMV (MCMV) M78 and the human herpesvirus (HHV) −6 and −7 protein U51 (13–16). The UL78 family are positionally conserved 7-transmembrane proteins that contain a DRL motif located within the second intracellular loop (ICL2) that is suspected to be necessary for G-protein coupling. UL78 family members undergo endocytosis from the cell surface like many GPCRs; however, only HHV U51 has been shown to bind chemokines and induce migration (17–20). UL78, R78, and M78 remain orphan GPCRs with no known ligands and recent structural analysis suggests that UL78 forms homotrimers that may occlude the putative ligand binding domain (21). A lack of UL78 expression does not impact HCMV lytic replication, however, R78 and M78 are necessary for efficient *in vitro* replication (16,22,23). RCMV R78 is expressed in many tissues and peripheral blood mononuclear cells in infected rats, and virus lacking R78 does not replicate in the spleen (15,24,25). MCMV M78 is important for transport of virus-infected cells to the salivary glands in part might be due to its ability to participate in the downregulation of MHC-II from the infected cell surface (26,27). In transient transfection assays, UL78 was shown to form heterodimers with the cellular chemokine receptors CXCR4 and CCR5 and reduce their cell surface expression as well as with HCMV vGPCR US28, which affected US28-mediated NF-κB activation, however the mechanism(s) for these findings has yet to be investigated (28,29). Thus, while there are some sequence similarities between the UL78 family members, they may functionally contribute to CMV infection in different ways.

Herein, we investigated the role of HCMV UL78 in latent infection of human embryonic stem cell (hESC) -derived CD34^+^ HPCs. Our results indicate that Gα_i_ coupling via the DRL motif of UL78 is essential for efficient reactivation from latent infection. To investigate the function of UL78, we performed proximity-dependent labelling experiments utilizing a recombinant virus expressing UL78 containing a C’ terminal fusion with TurboID in infected fibroblasts and during reactivation from latency in CD34^+^ HPCs. We identified a number of cellular and viral proteins as candidate UL78 interactors, including nuclear-localized proteins such as components of the nucleoporin complex, cellular and viral transcriptional regulators, and viral DNA replication machinery, suggesting that UL78 may localize to the nucleus, as has been observed for a number of cellular GPCRs (30–33). We determined that a fraction of both WT and G protein-coupling null mutant UL78 is detected at the nucleus using cell fractionation, luciferase assays, and immunofluorescent approaches. Together, our data indicate that UL78 coupling with Gα_i_ is essential for reactivation from latency and that UL78 localization to the nucleus suggests a novel function for this orphan HCMV GPCR.

## Results

### HCMV UL78 is Required for Efficient Viral Reactivation from Latent Infection

Several studies have shown that at least two of the G protein coupled-receptors encoded by HCMV are integral for establishing a latent infection and facilitating viral reactivation (8,11,34–36), however, the function of UL78 in this process is still unknown. To evaluate a potential role for UL78 in viral latency or reactivation, we utilized the TB40/E-GFP bacterial artificial chromosome (BAC) to generate a recombinant virus deficient in UL78 protein expression, but not gene expression. GalK-mediated recombination was used to place two contiguous stop codons immediately following the initiating methionine of UL78 (UL78-2XSTOP) (37). To determine the growth kinetics of the recombinant virus, both single and multistep growth kinetics were analyzed in primary human fibroblasts. Similar to previously published studies utilizing recombinant viruses lacking the entire UL78 open reading frame (ORF) (22,38), UL78-2XSTOP replicated with normal growth kinetics in this cell type **(Fig. S1).**

To determine whether UL78 has a role in the establishment of latent infection or the capacity of the virus to reactivate, we infected human embryonic stem cell (hESC)-derived CD34^+^ HPCs with either WT-HCMV (TB40/E-GFP) or UL78-2XSTOP (TB40/E-GFP-UL78-2XSTOP) viruses. At 48 hours post infection (hpi), viable, infected (GFP^+^), CD34^+^ HPCs were isolated via fluorescence activated cell sorting (FACS) and were seeded into long-term bone marrow culture (LTBMC) above a murine stromal support layer under conditions that favor latent infection, as previously described (9,39,40). After 12 days of LTBMC, half of the infected HPCs from each infection group were lysed by mechanical disruption to serve as a pre-reactivation control. The remaining intact HPCs and lysates were plated over monolayers of fibroblasts in reactivation supportive media supplemented with granulocyte-macrophage colony stimulating factor (GM-CSF) and granulocyte colony stimulating factor (G-CSF) to perform an extreme limiting dilution assay (ELDA) quantifying the frequency of infectious centers at three weeks post-plating (41). Comparable levels of infectious virus were present in lysed cells (pre-reactivation) infected with WT-HCMV or UL78-2XSTOP suggesting that UL78 has no effect on the establishment or maintenance of viral latency **(Fig. 1A, Fig. S2)**. In contrast, the frequency of infectious centers for the UL78-2XSTOP-infected cells did not increase in the presence of reactivation stimulus compared to WT-HCMV infected cells suggesting that UL78 is required for efficient reactivation from latency **(Fig 1A, Fig. S2)**. Since an observed deficit in viral reactivation can be caused by an inability to maintain viral genomes or genome-containing cells throughout latency, we quantified viral genome copies from infected CD34^+^ HPCs at the end of latent infection via quantitative PCR. A comparable number of viral genomes were present in cells infected with WT-HCMV or UL78-2XSTOP indicating that, despite the presence of viral DNA, viruses lacking UL78 are unable to produce infectious virions in HPCs stimulated to reactivate **(Fig. 1B)**. Together, these results indicate that UL78 plays an integral role in the viral reactivation process in CD34^+^ HPCs.

**Figure 1:**
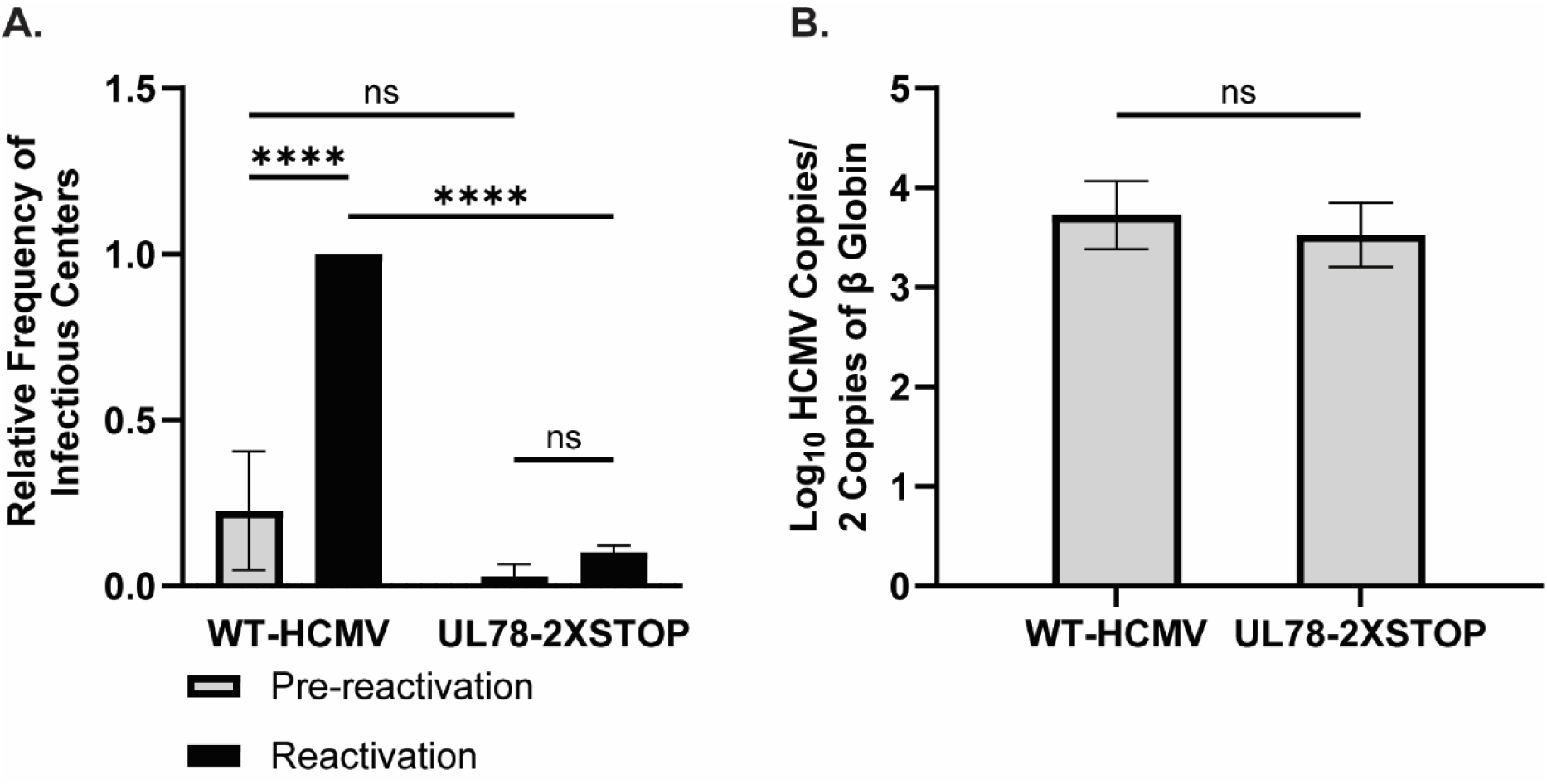
HCMV UL78 is Required for Reactivation from Latent Infection. hESC-derived CD34^+^ HPCs were infected with either WT-HCMV (TB40/E-GFP) or UL78-2XSTOP (TB40/E-GFP-UL78-2XSTOP) at a MOI of 2 for 48 hrs. Cells were FACS isolated for viable CD34^+^/GFP^+^ HPCs and were cultured above a murine stromal cell support layer for 12 days to establish latent infection. **(A)** At 14-dpi, half of the cells were treated with reactivation cocktail and plated onto a fibroblast monolayer (reactivation). The remaining cells did not receive treatment and were mechanically lysed (pre-reactivation). Reactivation was assessed by the frequency of infectious centers as determined via ELDA and compared to lysed cells (pre-reactivation) at 3 weeks post-plating. Data is shown as fold change in infectious centers, as compared to the WT-HCMV reactivation group, for triplicate experiments. Error bars represent the standard error of the mean. Statistical significance was calculated using two-way ANOVA followed by Tukey’s post-hoc analysis (*\*\*\*\*p < 0.0001*). **(B)** At 14-dpi, total DNA was harvested from infected CD34^+^ HPCs and viral genomes were quantified via qPCR using primers and probes specific for the viral UL141 gene. Viral genomes were normalized to total cell number using human β-globin as a reference gene. Data represents the mean Log_10_ transformed values for triplicate experiments. Error bars represent the standard error of the mean. Statistical significance was calculated using a student’s t-test.

### HCMV UL78 Coupling to Gα_i_ Heterotrimeric G-proteins via a Conserved DRL motif is Required for Reactivation from Latency

The DRY motif is a highly conserved sequence found in the second intracellular loop of most Class A GPCRs (42,43). Located at the boundary of transmembrane helix 3 (TM3) and intracellular loop 2 (ICL2), this motif forms an ionic lock to help maintain GPCR conformation and plays a crucial role in receptor activation. Specifically, the arginine residue within this motif stabilizes the receptor to facilitate G protein activation and subsequent signal transduction (44–46). UL78 contains a DRL motif that is conserved across the UL78 family. To identify the complement of G proteins that bind to UL78, we made use of a nLuc-based complementation assay measuring real-time interactions between receptors and heterotrimeric G protein complexes described previously (9,47). In this system, the C’ terminus of the GPCR is linked in frame to natural peptide (NP) while the Gα subunit is genetically fused to the complementing Large Bit (LgBiT). Proximity of the complementing fragments reconstitutes a functional luciferase protein whose activity can be measured with addition of substrate. To this end, we engineered an in-frame natural peptide-tag on the C’ terminus of the wild-type UL78 receptor. We utilized site directed mutagenesis to make alanine substitutions for the entire motif (DRL_133-135_ – AAA_133-135_) and the arginine specifically (DRL_133-135_ – DAL_133-135_) to serve as negative controls as these mutations would be predicted to affect G protein coupling. As a positive control, we used US28 as it has been shown to functionally couple to most Gα family members (9,48,49). Equivalent expression of each construct was verified by immunoblot in transiently transfected HEK-293 cells **(Fig. S3)**. In live cell GPCR interaction assays with LgBiT-Gα_i_, -Gα_q_, and -Gα_12_, the wildtype UL78 receptor exhibited similar Gα_i_ coupling to that of US28 **(Fig. 2A, D)**, but did not significantly couple to the other Gα isoforms **(Fig. 2B-D).** Moreover, both UL78 constructs containing mutations within the DRL motif showed significant attenuation of their ability to couple to the Gα_i_ family of G-proteins and did not show any increase in coupling breadth with LgBiT-Gα_q_ and -Gα_12_ **(Fig. 2)**. Together, these data indicate that UL78 Gα_i_-specific coupling requires the DRL motif.

**Figure 2:**
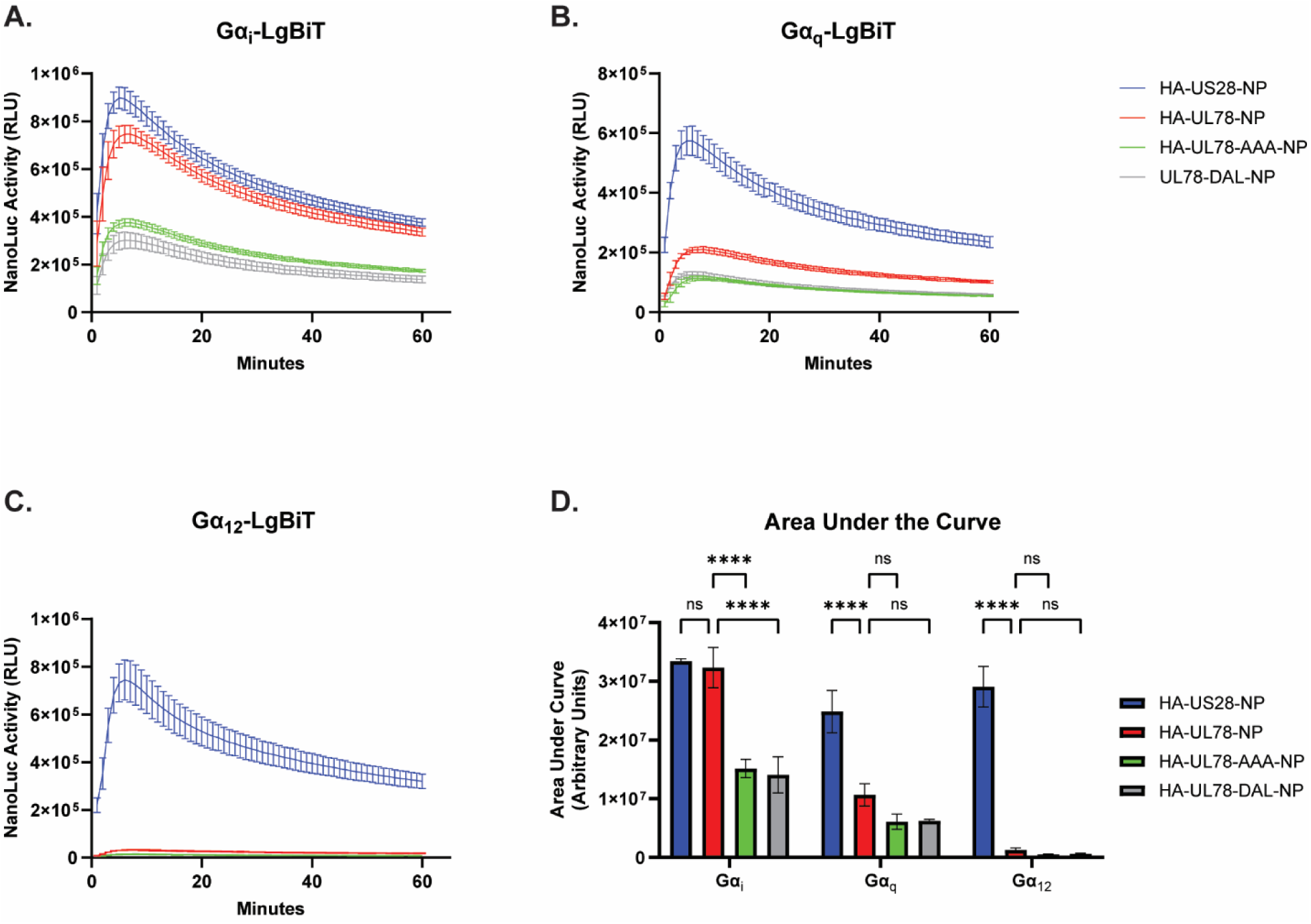
HCMV UL78 Preferentially Couples to Gα_i_ Isoforms via the DRL Motif. HEK-293 cells were transfected with the indicated constructs or the empty vector (EV) and LgBiT tagged **(A)** Gα_i_, **(B)** Gα_q_, or **(C)** Gα_12_. At 18 hrs post-transfection, media was exchanged with serum-free DMEM. At 6 hrs post-media replacement, luciferase activity was monitored for 60 min using the Nano-Glo Live Cell Assay System (Promega). Error bars represent the standard error of the mean between technical triplicates. **(D)** The area under the curve was calculated for each profile and plotted. Error bars represent the standard error of the mean between triplicate experiments. Statistical significance was calculated using two-way ANOVA followed by Dunnett’s multiple comparison post-hoc analysis *(****p < 0.001)*.

To assess the effects of UL78 G-protein coupling within the context of viral infection, we generated recombinant viruses using the TB40/E-GFP BAC and engineering the HiBiT tag onto the N’ terminus of UL78 (TB40/E-GFP-HB-UL78). An additional recombinant HiBiT tagged virus was generated containing an alanine substitution at position R134 (TB40/E-GFP-HB-UL78-DAL). Following reconstitution of these viruses, we evaluated their growth kinetics via single- and multi-step growth analyses in primary fibroblasts. Both recombinant viruses replicated with similar kinetics to each other, and to the parental TB40/E-GFP virus indicating that the addition of a HiBiT tag to UL78 and the substitution R134A have no effect on lytic replication **(Fig. S4)**. Furthermore, both recombinant viruses demonstrated immediate early, early, and late protein expression with similar kinetics **(Fig. 3A)**. UL78 protein expression was first observed at 24 hpi with peak expression observed between 72 – 96 hpi **(Fig 3A).** To determine whether the DRL motif mutations alter receptor expression at the cell surface, we conducted HiBiT surface vs. total expression assays in infected human fibroblasts (9). In this assay, the total luminescence emitted by HiBiT tagged UL78 in lysed cells is compared with that of live cells where only surface UL78 can interact with the complementing LgBiT. The ratio calculated between the surface and total luminescence indicates the fraction of UL78 that is present at the surface. Consistent with previously published data (19,20), we observed that the majority of UL78 localized intracellularly **(Fig. 3B)**. Additionally, no appreciable difference in surface expression was detected between the two recombinant viruses indicating that mutation within the DRL motif of UL78 does not significantly affect cellular surface expression **(Fig. 3B)**.

**Figure 3:**
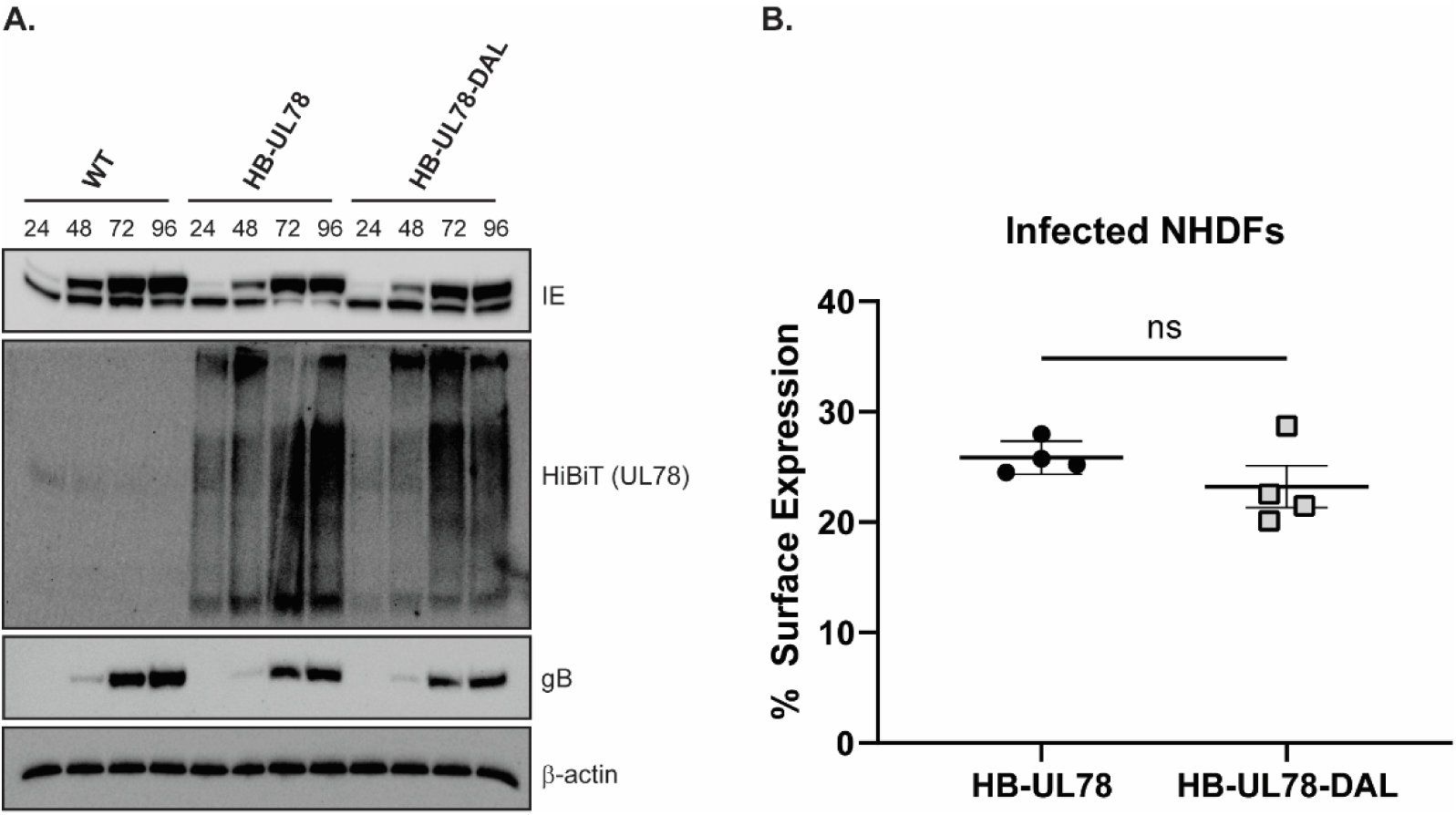
HCMV UL78 Expression and Membrane Localization. **(A)** NHDFs were infected with the indicated viruses at an MOI of 2 and lysates were harvested at the specified time points post-infection. The presence of the indicated proteins was determined by immunoblot using the indicated antibodies or the Nano-Glo HiBiT Blotting System (Promega). **(B)** NHDFs were infected with the indicated HiBiT-tagged viruses at an MOI of 2. At 72-hpi, surface expression was measured using the Nano-Glo HiBiT Extracellular and Lytic detection Systems (Promega). Error bars represent the standard error of the mean between triplicate experiments. Statistical significance was calculated using one-way ANOVA followed by Dunnett’s multiple comparison post-hoc analysis.

To determine whether the observed reactivation deficit with recombinant viruses lacking UL78 protein expression **(Fig. 1A)** can be recapitulated with a virus deficient in UL78 Gα_i_ protein coupling, we infected hESC-derived CD34^+^ HPCs with either WT-HCMV (TB40/E-GFP) or HB-UL78-DAL (TB40/E-GFP-HB-UL78-DAL). Viable, GFP^+^, CD34^+^ HPCs were isolated via FACS and after 12 days of LTBMC, both cells stimulated to reactivate and lysates from unstimulated cells were plated onto fibroblast monolayers to assess infectious center frequency. When compared to infection with WT-HCMV, cells infected with HB-UL78-DAL exhibited major deficits in the ability of the virus to efficiently reactivate from latent infection **(Fig. 4A, Fig. S5)**. Viral genome copies from infected CD34^+^ HPCs at the end of LTBMC were comparable in cells infected with WT-HCMV and HB-UL78-DAL suggesting that viral genomes, or genome-containing cells, were not lost over the latency culture period **(Fig. 4B)**. Collectively, these results indicate that UL78 – Gα_i_ coupling is not required for the establishment of viral latency in CD34^+^ HPCs but is required for efficient reactivation from latent infection.

**Figure 4:**
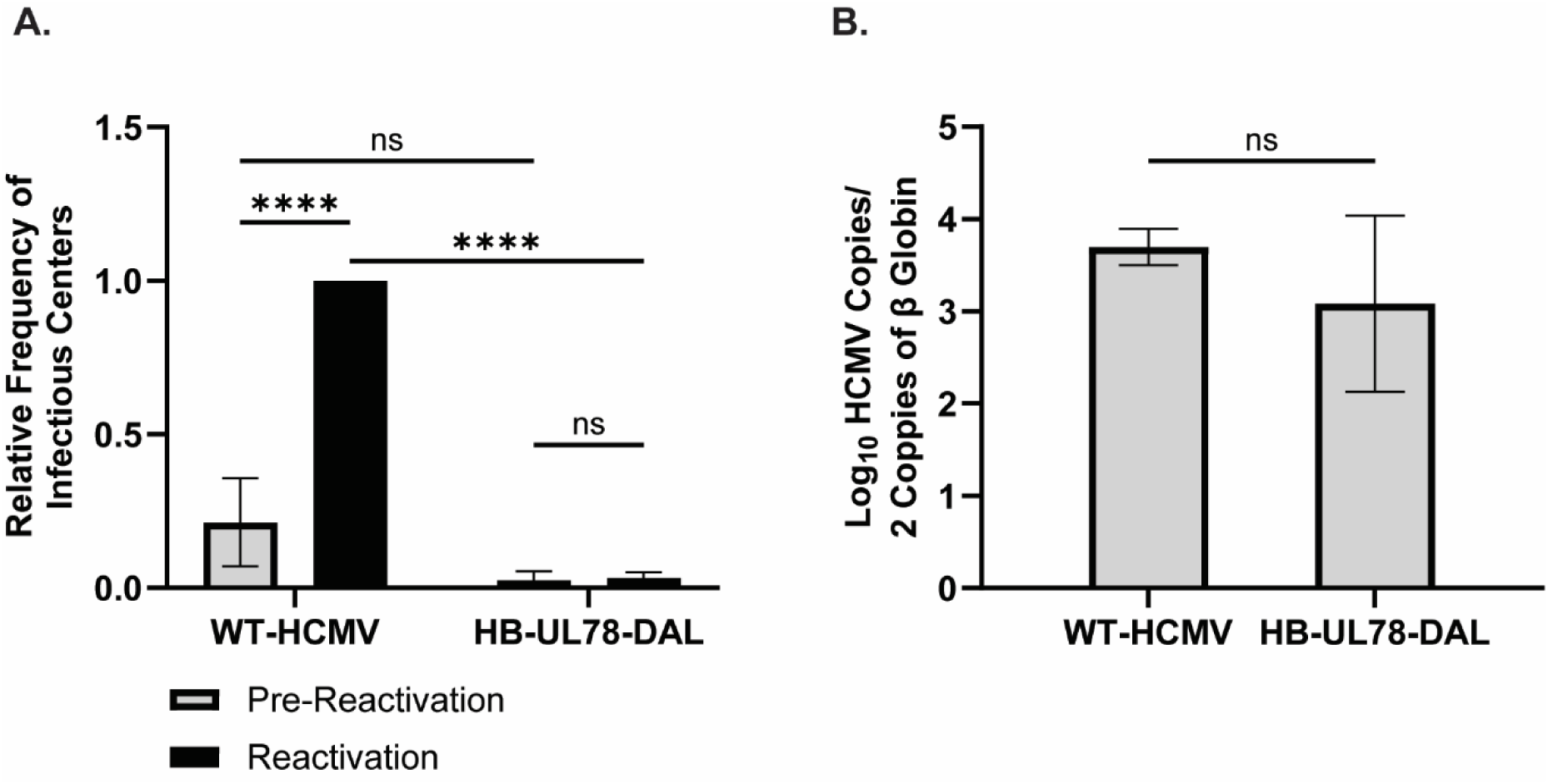
Attenuating HCMV UL78 G-Protein Coupling Results in Viral Reactivation Deficits. hESC-derived CD34^+^ HPCs were infected with either WT-HCMV (TB40/E-GFP) or HB-UL78-DAL (TB40/E-GFP-HB-UL78-DAL) at a MOI of 2 for 48 hrs. Cells were FACS isolated for viable CD34^+^/GFP^+^ HPCs and were cultured above a murine stromal cell support layer for 12 days to establish latent infection. **(A)** At 14-dpi, half of the cells were treated with reactivation cocktail and plated onto a fibroblast monolayer (reactivation). The remaining cells did not receive treatment and were mechanically lysed (pre-reactivation). Reactivation was assessed by the frequency of infectious centers as determined via ELDA and compared to lysed cells (pre-reactivation) at 3 weeks post-plating. Data is shown as fold change in infectious centers, as compared to the WT-HCMV reactivation group, for triplicate experiments. Error bars represent the standard error of the mean. Statistical significance was calculated using two-way ANOVA followed by Tukey’s post-hoc analysis (*\*\*\*\*p < 0.0001*). **(B)** At 14-dpi, total DNA was harvested from infected CD34^+^ HPCs and viral genomes were quantified via qPCR using primers and probes specific for the viral UL141 gene. Viral genomes were normalized to total cell number using human β-globin as a reference gene. Data represents the mean Log_10_ transformed values for triplicate experiments. Error bars represent the standard error of the mean. Statistical significance was calculated using a student’s t-test.

### Identification of the HCMV UL78 Interactome During Lytic Infection

A limited number of previous studies have shown that UL78 forms heterodimeric interactions with host (CXCR4 and CCR5) and viral (US28) chemokine receptors to impair or augment surface expression and downstream signaling activity (28,29). While valuable, these overexpression models monitored interactions in transiently transfected cells, which do not recapitulate the conditions of viral infection. Furthermore, UL78 may exhibit additional cell type specific interactions with host and viral proteins to modulate signal transduction that have yet to be captured. To characterize the UL78 interactome during viral infection, a recombinant HCMV was generated containing the biotin ligase TurboID (50) linked in-frame to the C’ terminal tail of UL78 (TB40/E-GFP-UL78-TurboID). The recombinant virus replicated with similar growth kinetics in primary fibroblasts relative to the parental virus indicating that the addition of the TurboID enzyme to UL78 has no effect on lytic replication **(Fig. S6)**. To identify the UL78 interactome, fibroblasts were mock infected or infected with WT-HCMV (TB40/E-GFP) or UL78-TurboID (TB40/E-GFP-UL78-TurboID). At 72 hpi, exogenous biotin was added to the culture media for an additional six hours. Cellular lysates were harvested, and the resulting biotin-conjugated proteins were purified via streptavidin-mediated bead-based precipitation. Efficient labeling and purification were verified by immunoblot probing for HRP-conjugated streptavidin **(Fig. 5A)**. The identity and relative abundance of the purified biotin-conjugated proteins were determined by label-free quantitative (LFQ) liquid chromatography tandem mass spectrometry (LC-MS/MS). After excluding proteins identified as potential contaminants by comparison against the CRAPome data repository (51), our analysis revealed 1138 host and 32 viral proteins that showed a significant level of enrichment within the dataset **(Fig. 5B,C, Data S1)**. Interestingly, neither of the previously identified host chemokine receptors reported to interact with UL78 (e.g., CXCR4 and CCR5) (29) were enriched within the dataset, however, the previously identified viral candidate interaction partner (US28) was identified above the significance threshold (28,34). Consistent with a recently published study (21), we identified several Gα_i_ isoforms, but not other G protein families, as candidate interaction partners of UL78 **(Fig. 5B, Data S1)**, validating the results of our split-nano luciferase assays **(Fig. 3)**. Reactome over-representation analysis of the enriched host proteins in proximity to UL78 revealed several cellular processes relating to trafficking, signal transduction, cytoskeletal remodeling, and processes related to nuclear import **(Fig. S7A, Data S2)**. Surprisingly, we identified many nuclear-localized cellular proteins as candidate UL78 interactors **(Figs. 5B, Data S1,2)**, including many of the components of the nuclear pore complex, nuclear membrane proteins, and transcription factors. Additionally, importin, Rab, SNX, and RanGDP proteins were found in proximity to UL78, suggesting a mechanism for translocation to the nucleus. Furthermore, viral proteins, including components of the DNA replication machinery, transcriptional regulators, and proteins that regulate nuclear egress, are candidate interaction partners for UL78 **(Fig. 5C, Data S1)**. To confirm the candidate UL78 nuclear interaction partners during lytic infection, we performed a second proximity-dependent labeling experiment in which NHDF cells were infected and treated with biotin as previously described, and then cytosolic and nuclear fractions were separated prior to tagged protein purification. Efficient labeling and fractionation were verified by immunoblot prior to LC-MS/MS analysis where we observed visible labeling within the nuclear fraction of cell lysates **(Fig. 5D)**. Quantitative mass-spectrometry and downstream analysis revealed 1592 host and 83 viral proteins as candidate interaction partners of UL78 within the nuclear fraction **(Fig. 5E, F, Data S3)**. Similar to previous experiments, we identified several transcription factors, members of the nuclear pore complex, and Gα_i_ isoforms **(Fig. 5E, Data S3)**. Reactome over-representation analysis of host candidate interaction partners identified several previously identified processes such as nuclear import, trafficking, and transcription as well as additional processes involved with RNA biogenesis and chromatin remodeling **(Fig. S7B, Data S4)**. Moreover, many of the viral proteins enriched within this dataset are known nuclear proteins involved in viral DNA replication, transcription, DNA packaging, encapsidation, and nuclear egress **(Fig. 5F, Data S3)**. Taken together, our interactome analysis demonstrates that UL78 uniquely interacts with nuclear host and viral proteins during lytic infection conditions and offers novel insight into the function and signal transduction capability of UL78.

**Figure 5:**
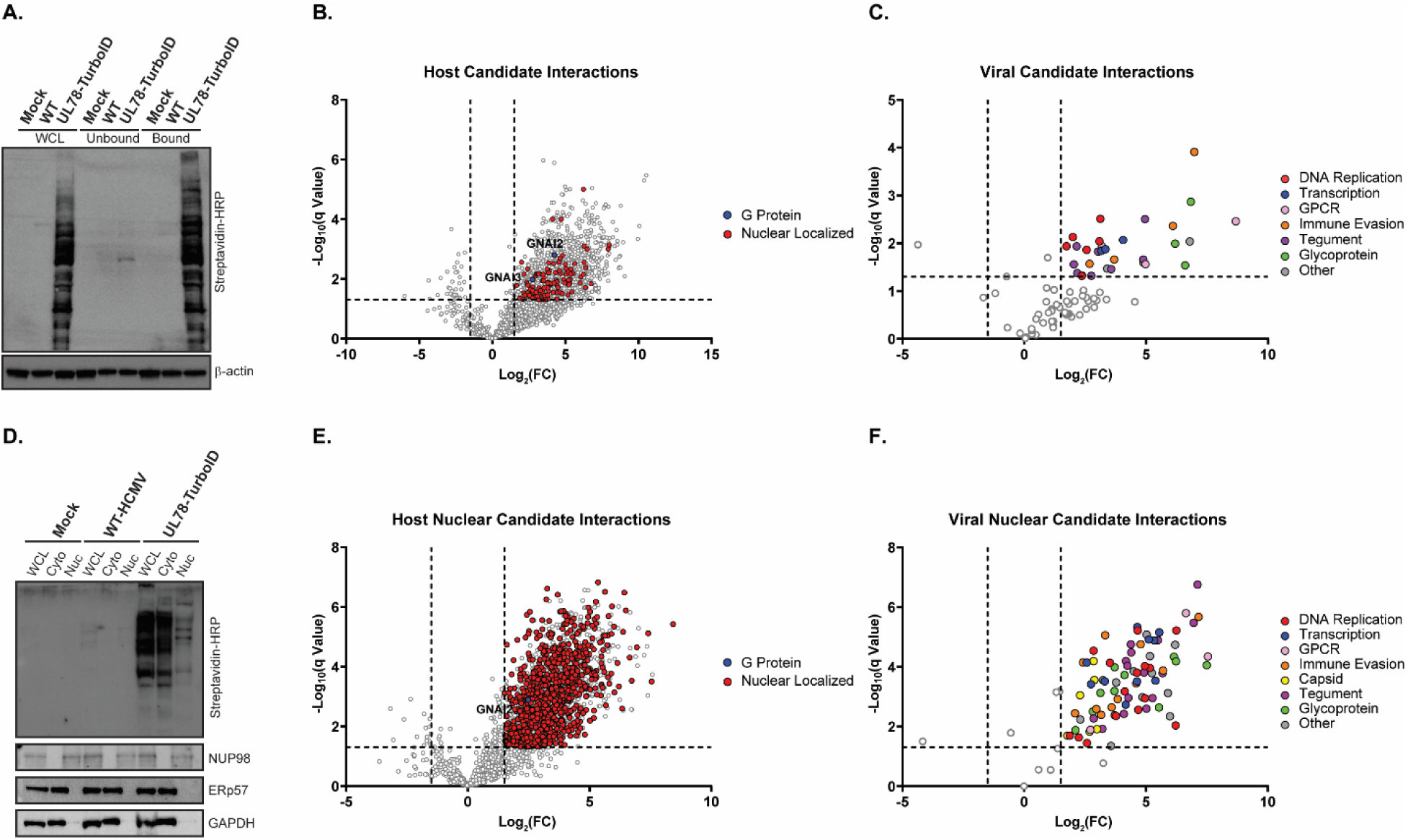
Interactome Analysis of HCMV UL78. **(A)** NHDF cells were infected with either WT-HCMV (TB40/E-GFP) or UL78-TurboID (TB40/E-GFP-UL78-TurboID) at an MOI of 2 or mock infected. At 72-hpi, the cell culture medium was supplemented with biotin (50 ug/mL) for 6 hrs. Tagged protein were bound to neutravidin beads and were incubated overnight prior to purification. **(A)** Efficient labeling and purification were evaluated by immunoblot using the indicated antibodies on whole cell lysates, unbound fraction, and bound fraction. Purified proteins were subjected to quantitative LC-MS/MS analysis. Representative blot shown for triplicate experiments. **(B)** Volcano plot of host candidate interaction partners of UL78. **(C)** Volcano plot of viral candidate interaction partners of UL78. **(D)** NHDF cells were infected and treated under the same conditions as previously described. At 72-hpi, lysates were harvested and were fractionated using the NE-PER extraction kit (ThermoFisher Scientific). Efficient labeling and purification were evaluated by immunoblot using the indicated antibodies on whole cell lysates, cytoplasmic fraction, and nuclear fraction. Purified proteins were subjected to quantitative LC-MS/MS analysis. Representative blot shown for triplicate experiments. **(E)** Volcano plot of nuclear enriched host candidate interaction partners of UL78. **(F)** Volcano plot of nuclear enriched viral candidate interaction partners of UL78.

### Examining HCMV UL78 Interactions During Viral Reactivation

Since UL78 is important for reactivation from latency, we wanted to identify host and viral interaction partners of UL78 in the context of HCMV reactivation. To this end, we conducted proximity-dependent labeling experiments in CD34^+^ HPCs stimulated to reactivate from latent infection. In this experiment, hESC-derived CD34^+^ HPCs were infected with either WT-HCMV (TB40/E-GFP) or UL78-TurboID (TB40/E-GFP-UL78-TurboID) and were cultured in the same manner as the above experiments **(Fig. 6A)**. After 12 days of LTBMC, cells were plated in transwells above a fibroblast monolayer in reactivation supportive media supplemented with exogeneous biotin and incubated for an additional 16 hours prior to cell lysis. Efficient labeling of proteins in proximity of UL78 was confirmed on whole cell lysates via immunoblot probing with HRP-conjugated streptavidin prior to purification and tryptic digestion **(Fig. 6B)**. The resultant peptides were subjected to LC-MS/MS analysis. After contaminant filtering, we identified 1075 host and 61 viral proteins as candidate interaction partners of UL78 **(Data S5)**. Notably, we again identified Gα_i_ as the only Gα isoform present within this dataset. Additionally, we identified the other viral GPCRs (UL33, US27, and US28), immune evasion proteins (UL40, US23, and US26), and transcriptional activators (UL49, UL69, UL82, and UL122). When compared to our previous proximity-dependent labeling experiments conducted in fibroblasts under lytic conditions **(Fig. 5, S6)**, a total of 354 common and 721 unique host hits were identified in CD34^+^ HPCs **(Fig. 6C)**. In a similar manner, this dataset contained 54 common and 9 unique viral proteins **(Fig. 6D**, **Table 1)**. Finally, Reactome over-representation analysis of the identified host proteins revealed enrichment in several processes related to RNA processing, trafficking, protein post-translational modifications, and members of the nuclear pore complex **(Fig. 6E, Data S6)**. Together, these results confirm the findings from our previous proximity-dependent labeling experiments in a reactivation model and suggest novel interaction partners and nuclear localization, which will both aid in deciphering the function of UL78 during HCMV pathogenesis.

**Figure 6:**
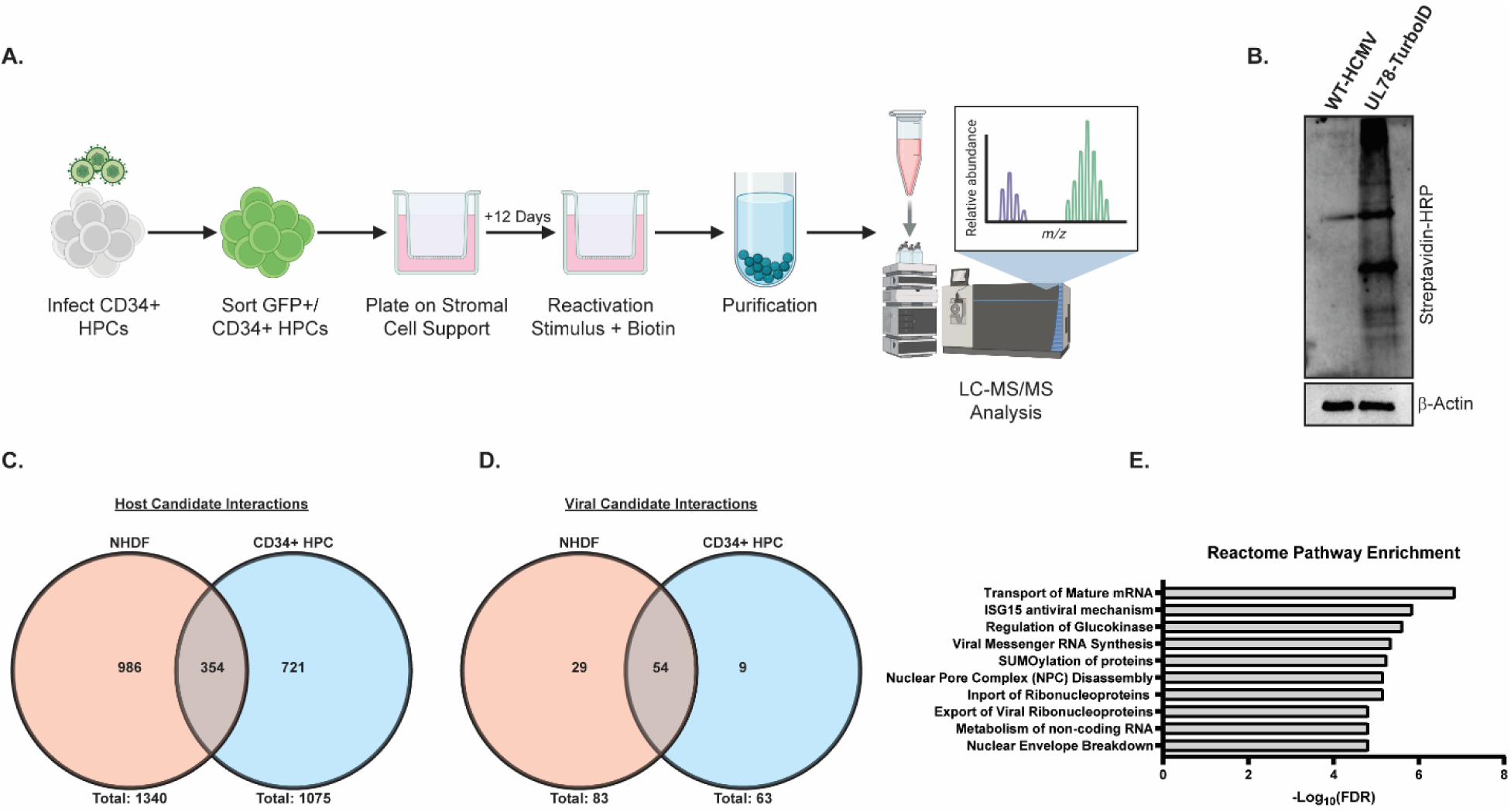
Comparative Analysis of the HCMV UL78 Interactome During Viral Reactivation. **(A)** hESC-derived CD34^+^ HPCs were infected with either WT-HCMV (TB40/E-GFP) or UL78-TurboID (TB40/E-GFP-UL78-TurboID) at a MOI of 2 for 48 hrs. Cells were FACS isolated for viable CD34^+^/GFP^+^ HPCs and were cultured above a murine stromal cell support layer for 12 days to establish latent infection. At 14-dpi, latently infected cells were plated above a fibroblast monolayer and treated with reactivation cocktail supplemented with biotin (50 ug/mL) for 16 hrs. Biotinylated proteins were purified and were subjected to quantitative LC-MS/MS analysis. **(B)** Efficient labeling was evaluated by immunoblot using the indicated antibodies on whole cell lysates. **(C)** Candidate host UL78 interaction partners were compared between infected NHDFs and CD34^+^ HPCs. **(E)** Candidate viral UL78 interaction partners were compared between infected NHDFs and CD34^+^ HPCs. **(E)** Reactome over-representation analysis of high confidence candidate interaction partners.

**Table 1:** Viral Proteins Within the UL78 Interactome in Infected Fibroblasts and CD34^+^ hematopoietic progenitor cells.

| Gene Symbol | Fibroblast Lytic Infection |  | CD34<sup>+</sup> HPC Reactivation | Function |
| --- | --- | --- | --- | --- |
|  | Log<sub>10</sub>(q Value) | Log<sub>2</sub>(FC) | Identified |  |
| RL11 | 1.592 | 4.898 | Peak Found | IgG Fc Binding |
| RL12 | N/A | N/A | High | IgG Fc Binding |
| **UL13** | 1.660 | 4.884 | Peak Found | Unknown |
| **UL24** | N/A | N/A | High | Immune evasion |
| **UL26** | N/A | N/A | High | MIEP activator |
| **UL31** | N/A | N/A | High | Nucleolar organization |
| **UL32** | N/A | N/A | High | Tegument protein, viral DUB main target |
| **UL34** | N/A | N/A | High | Transcriptional repressor |
| **UL35** | N/A | N/A | High | Tegument protein, viral IE gene expression |
| **UL36** | N/A | N/A | High | Anti-apoptotic, inhibits caspase 8 activation |
| **UL40** | 2.362 | 6.098 | Peak Found | NK cell evasion |
| **UL45** | 1.376 | 2.172 | High | Immune evasion, blocks NF-kB signaling |
| **UL46** | N/A | N/A | High | Capsid protein |
| **UL47** | 1.320 | 2.754 | N/A | Tegument protein, viral DNA release from capsid |
| **UL48** | 1.937 | 2.153 | High | Immune evasion, deubiquitin protease |
| **UL49** | 1.842 | 3.164 | Peak Found | Late viral gene expression, DNA replication |
| **UL50** | 1.828 | 3.030 | High | Nuclear egress, disruption of nuclear lamina |
| **UL52** | 2.511 | 3.124 | High | DNA cleavage, packaging |
| **UL53** | N/A | N/A | High | Nuclear egress, disruption of nuclear lamina |
| **UL54** | 2.042 | 3.084 | N/A | DNA polymerase |
| **UL56** | 1.860 | 2.560 | Peak Found | Terminase subunit |
| **UL57** | N/A | N/A | High | ssDNA binding protein |
| **UL69** | 1.875 | 3.329 | High | mRNA nuclear export, cell cycle block |
| **UL70** | 2.130 | 1.992 | Peak Found | Primase, DNA synthesis |
| **UL71** | 2.506 | 4.957 | High | Virus spread and release |
| **UL72** | 1.457 | 3.566 | Peak Found | Transcription-replication machinery |
| **UL78** | 2.459 | 8.683 | Peak Found | vGPCR |
| **UL80** | N/A | N/A | High | Assembly Protein |
| **UL82** | N/A | N/A | High | pp71, MIEP activator, binds Rb |
| **UL84** | 1.939 | 1.729 | High | UTPase activity, suppresses transcription of IE2 |
| **UL85** | N/A | N/A | High | Minor capsid protein |
| **UL89** | N/A | N/A | High | Terminase ATPase subunit, DNA cleavage |
| **UL95** | N/A | N/A | High | Late viral gene expression |
| **UL96** | N/A | N/A | High | Nucleocapsid stability |
| **UL97** | N/A | N/A | High | Ser-thr kinase, regulates nuclear export, DNA replication |
| **UL98** | N/A | N/A | High | Alkaline nuclease |
| **UL100** | 1.538 | 6.603 | N/A | gM, recruits Rab11 |
| **UL102** | N/A | N/A | High | DNA helicase-primase |
| **UL112** | N/A | N/A | High | Transcriptional activator |
| **UL114** | 1.319 | 2.362 | N/A | Uracil DNA glycosylase, DNA replication |
| **UL121** | 1.993 | 6.190 | N/A | Unknown |
| **UL122** | 2.068 | 4.053 | High | IE2, negative regulator of MIEP |
| **UL132** | 5.039 | 6.836 | High | Viral glycoprotein |
| **UL138** | 1.661 | 3.690 | N/A | Silences IE1 transcription, latency |
| **UL140** | N/A | N/A | High | Unknown |
| **UL148** | N/A | N/A | High | NK cell evasion |
| **UL150** | 1.660 | 4.890 | N/A | Unknown |
| **US8** | 1.326 | 2.368 | N/A | Antagonize TLR signaling |
| **US12** | 1.470 | 3.402 | N/A | NK cell evasion |
| **US22** | 1.559 | 2.032 | High | Unknown |
| **US23** | 1.569 | 2.683 | Peak Found | Unknown |
| **US26** | 3.975 | 6.977 | Peak Found | Unknown |
| **US27** | 1.562 | 4.977 | Peak Found | vGPCR |
| **US28** | 1.941 | 3.181 | Peak Found | vGPCR, chemokine receptor |
| **US30** | 2.038 | 6.791 | N/A | Unknown |
| **TRS1** | N/A | N/A | High | Binds UL44, inhibits PKR |

### HCMV UL78 Localizes to the Nucleus During Infection

A limited number of cellular GPCRs have been detected in the nuclear envelope and nucleus, and have been shown to play an important role in host signaling and cell cycle regulation (30–32). Since UL78 proximity-dependent labelling experiments identified several host and viral nuclear-localized proteins during infection, we next used orthogonal methods to validate whether UL78 is present at the nucleus. Previous studies using transient overexpression and infection models clearly detect UL78 at the cell surface and in cytoplasmic endocytic vesicles (19,20). In addition, **Figure 4B** demonstrates cell surface expression of a fraction of UL78. To assess whether UL78 also localizes to the nucleus, we infected fibroblasts with WT-HCMV (TB40/E-GFP), HB-UL78 (TB40/E-GFP-HB-UL78), and HB-UL78-DAL (TB40/E-GFP-HB-UL78-DAL) viruses for 72 hours and performed cell fractionation to separate cytoplasmic and nuclear fractions. As shown by immunoblots from fractionated lysates, HB-UL78 and HB-UL78-DAL were detected in both the cytoplasmic and nuclear fractions, suggesting that UL78 localizes to the nucleus during lytic infection and that G protein coupling may not be required for translocation or retention at the nucleus **(Fig. 7A)**. To confirm the nuclear localization of UL78, we performed HiBiT split luciferase assays on fractionated lysates. In these experiments, we utilized WT-HCMV and a recombinant virus with a HiBiT tag on the N’ terminus of the membrane protein UL8 (HB-UL8) as negative controls (52). Similar to observations from immunoblot experiments **(Fig. 7A)**, we detected both HB-UL78 and HB-UL78-DAL within the nuclear fraction of these cell lysates **(Fig. 7B)**. Moreover, the signal in nuclear extracts obtained from cells infected with virus expressing HB-UL8 was negligible when compared to the background readings from cells infected with WT-HCMV **(Fig. 7B).** We further validated these findings using immunofluorescence in the context of fibroblast infection. WT-HCMV, HB-UL78, and HB-UL78-DAL viruses were used to infect human fibroblasts for 72 hours and UL78 localization was detected by confocal microscopy using an antibody to HiBiT. During HB-UL78 infection, UL78 could be detected in punctate structures consistent with previous reports showing localization in intracellular vesicles. However, HiBiT signal could also be detected at the nuclear membrane of HB-UL78 and HB-UL78-DAL-infected cells **(Fig. 8A, B)**, further supporting that a fraction of UL78 is nuclear and that this localization may be independent of G-protein coupling. Together, these data strongly support the nuclear localization of a portion of UL78 and suggest novel intra-nuclear functions for the GPCR.

**Figure 7:**
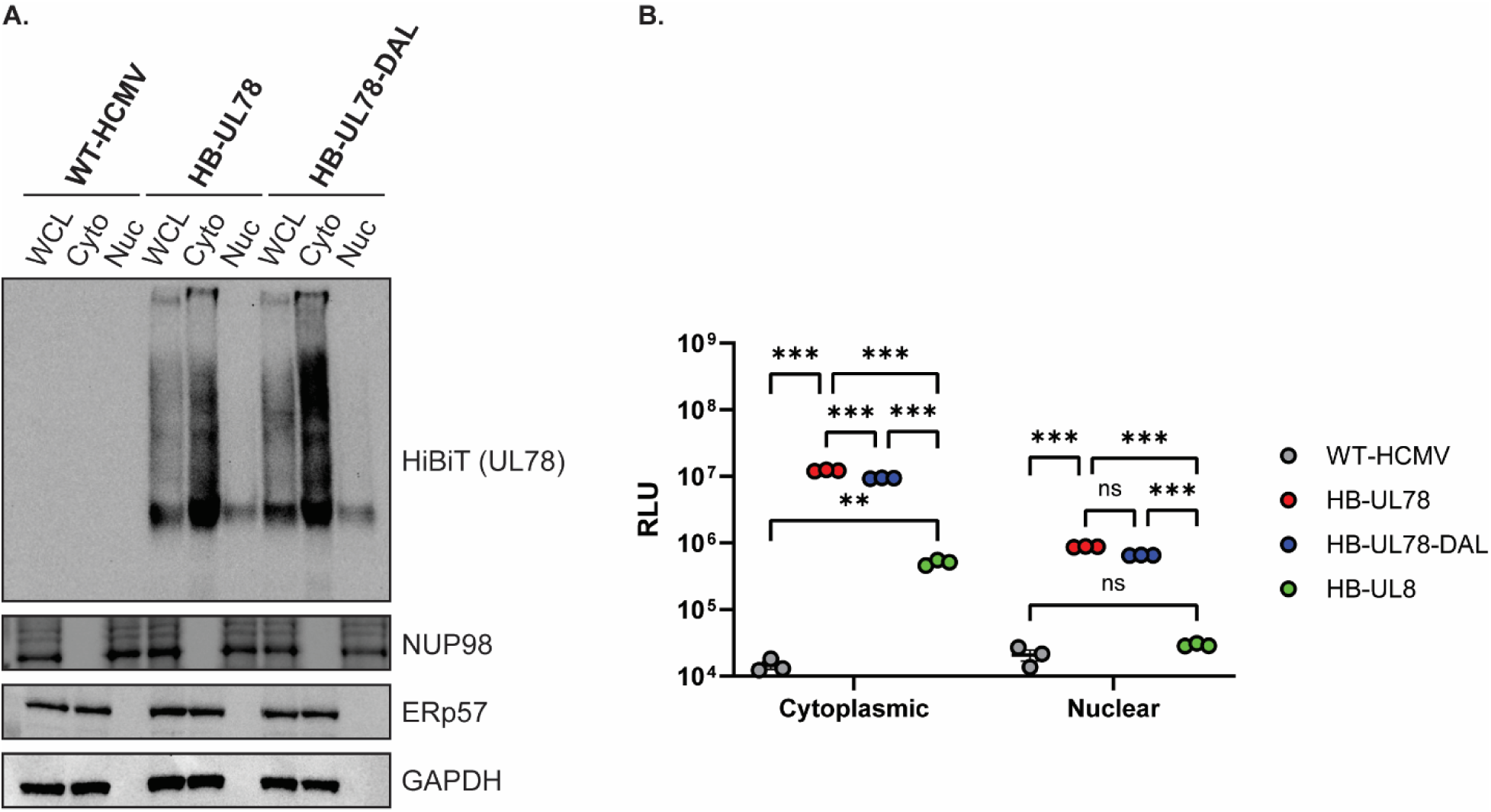
A Subset of HCMV UL78 Is Targeted to the Nuclear Envelope During Lytic Replication. **(A)** NHDFs were infected with the indicated viruses at an MOI of 2. At 72-hpi, whole cell lysates were harvested and were fractionated using the NE-PER extraction kit (ThermoFisher Scientific). The presence of the indicated proteins was determined by immunoblot using the indicated antibodies or the Nano-Glo HiBiT Blotting System (Promega). Representative blot shown from triplicate experiments. **(B)** NHDFs were infected with the indicated viruses at an MOI of 2. At 72-hpi, whole cell lysates were harvested and were fractionated using the NE-PER extraction kit (ThermoFisher Scientific). The presence of the indicated proteins was determined using the Nano-Glo HiBiT Lytic detection Systems (Promega). Error bars represent the standard error of the mean between triplicate experiments. Statistical significance was calculated using two-way ANOVA followed by Dunnett’s multiple comparison post-hoc analysis *(**p < 0.01, ***p < 0.005)*.

**Figure 8:**
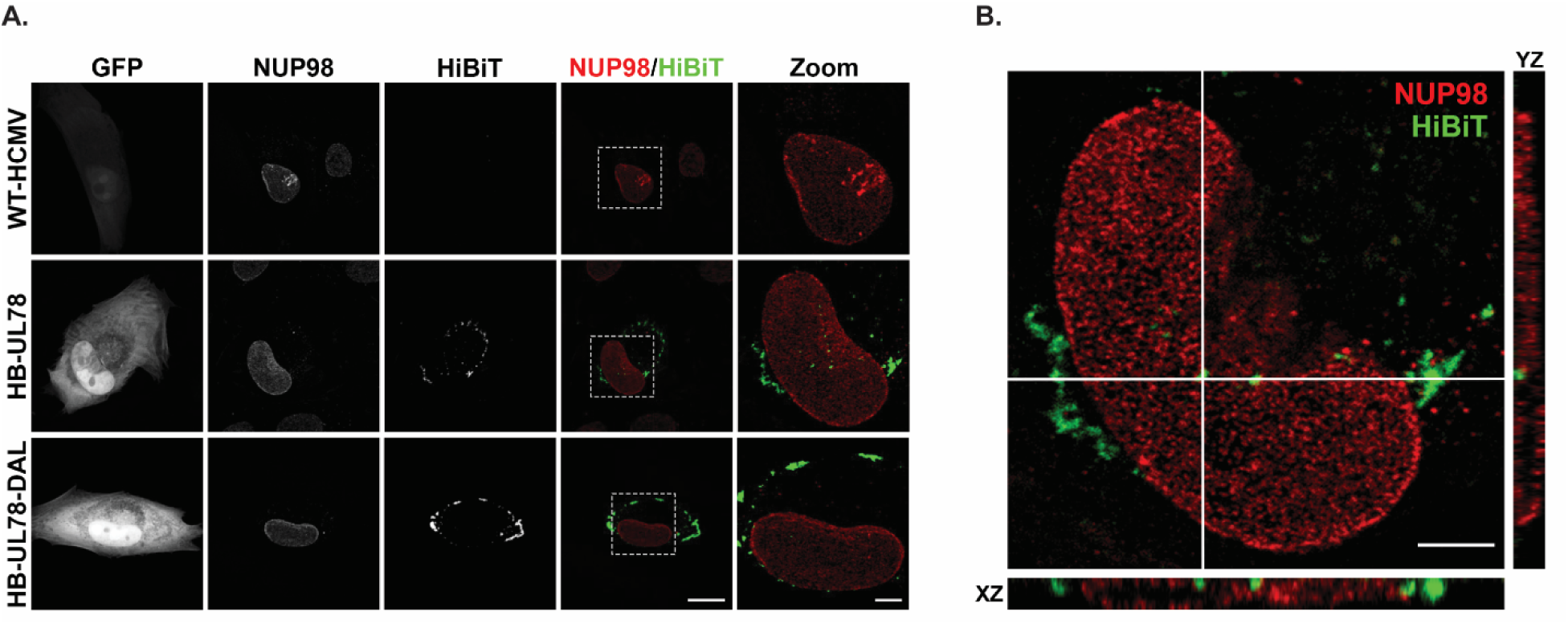
A Fraction of HCMV UL78 Localizes to the Nuclear Envelope During Lytic Infection. NHDF cells were plated on glass coverslips and infected at an MOI of 0.5 with the indicated viruses. Cells were fixed 72-hpi and stained for NUP98 and HiBiT. **(A)** Representative images are shown as the maximum projection from a z-stack. Right panels show overlay of NUP98 (red) and HiBiT (green). White dotted lines indicate region of interest used for magnified images (right panel). Scale bar, 20µm (four left panels) or 5µM (right panel). **(B)** A single z-stack of the HB-UL78-infected cell shown in panel A, along with orthogonal views in XY and XZ planes for the area of interest (indicated by white lines). Scale bar, 5µM.

## Discussion

A mechanistic role for the HCMV-encoded G protein coupled receptor UL78 in viral pathogenesis has remained elusive. In this study, we demonstrate that UL78 joins the viral genes UL7, UL8, UL33, UL81-82ast (LUNA), UL135, UL136, and US28 as well as viral miRNAs (miR-UL36, miR-UL112, and miR-UL148D) as factors required for HCMV reactivation that do not impact viral latency establishment or maintenance in myeloid lineage cells (8,9,11,34,39,52–56). Herein, we establish that recombinant HCMV that lacks UL78 protein expression or contains a single amino acid substitution in the DRL motif (DAL) were both unable to reactivate from latency in CD34^+^ HPCs. We also demonstrate that while UL78 specifically couples to Gα_i_ heterotrimeric G proteins, the UL78 DAL mutant failed to couple, which is consistent with the role of this motif in G protein coupling for other GPCRs. Combined, these findings suggest that UL78 coupling to Gα_i_ is necessary to promote viral reactivation. Gα_i_ specificity was also observed in UL78-proximity labeling where only Gα_i_ family members were identified as enriched in both HCMV infected human fibroblasts and latently infected CD34^+^ HPCs undergoing the early stages of viral reactivation. Moreover, an important, and yet unexpected, finding obtained through analysis of the proximity labeling experiments in both cell types positioned UL78 at the nucleus and identified a potential pathway of nuclear translocation for the viral GPCR (**Fig. 9**).

**Figure 9:**
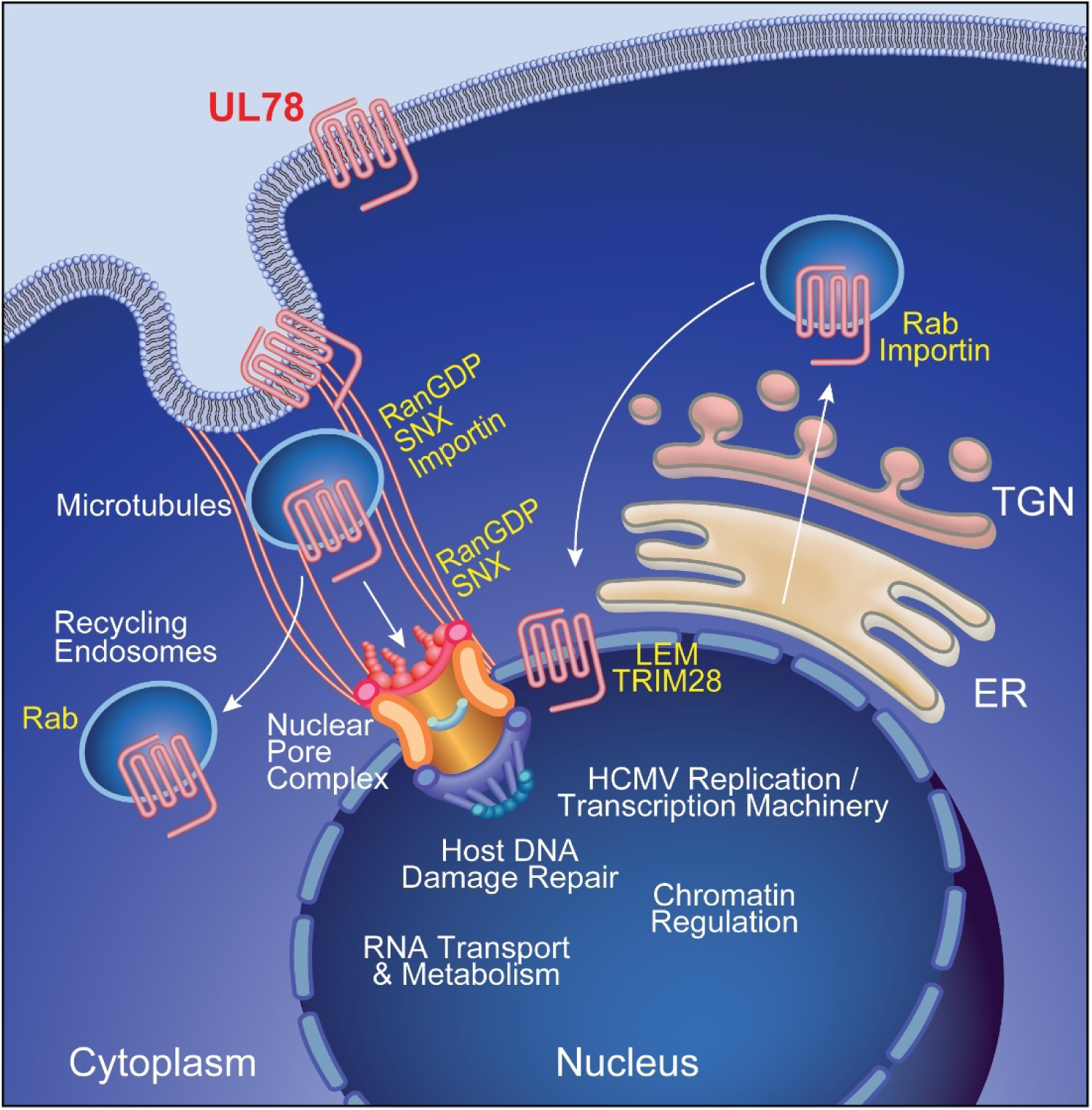
Proposed Model of HCMV UL78 Nuclear Translocation. HCMV UL78 translocates to nucleus via either agonist-dependent (RanGDP, SNX, Importin) or -independent (Rabs, Importin) mechanisms. Once at the host cell nucleus, HCMV UL78 is able to facilitate efficient viral reactivation, in a G protein coupling-dependent manner, likely acting as a scaffold to help regulate viral genome accessibility and promote viral transcription during reactivation.

While the role of UL78 and requirement of G protein coupling in the viral reactivation process are both clear, further characterization is needed to determine whether nuclear localization, signaling, and/or interactions with host and viral machinery located in the nucleus are required to promote reactivation. A functional role for nuclear GPCRs has become better appreciated in recent years (57,58). Some cellular GPCRs translocate from the plasma membrane to the nucleus upon ligand binding and activation, whereas others appear resident at the nucleus (59,60). Nuclear-localized GPCRs can activate ‘classical’ signaling cascades within the nucleus as G-proteins, GRKs, β-arrestin and components of many signal transduction pathways are readily detected in the nucleus (61,62) and nuclear GPCR activity can induce the phosphorylation of signaling intermediates as well as Ca^2+^ and cAMP flux (63). Additionally, some cellular GPCRs interact directly with transcriptional regulators and/or host DNA to impact gene expression. For example, nuclear translocated F2rl1 (PAR2) interacts with the transcription factor Sp1 and enhances expression of the *Vegfa* gene leading to neovascularization during mouse retinal development (64). Intriguingly, this study demonstrated that nuclear-localized PAR2 activated different transcriptional responses from plasma membrane localized PAR2 suggesting a dichotomy of cellular outcomes based on GPCR location.

Based upon our reactivation and proximity labeling data, we hypothesize that UL78 localized at or near the nuclear pore complex (NPC) is acting as a scaffold to help regulate viral genome accessibility and promote viral transcription during reactivation. This hypothesis is well founded in that the NPC plays a critical role in regulating host chromatin state and gene transcription by recruiting and organizing histone regulatory/remodeling complexes as well as by positioning the open chromatin near the NPC opening to facilitate easy access to transcription factors and nucleotide pools (65–68). The UL78 proximity data was enriched for members of the NPC including NUPs (35, 42, 50, 62, 88, 98, 133, 153, 155, 188, 214, 358), transcription factors (ATF6, JUN B, RelA, GTFIIF, SUMO1, YAP1), RNA metabolism (GTF3C1, POLII, SF1, SYMPK), as well as chromatin remodeling proteins (AHCTF1, LEM, LBR, SAP18, TRIM25, TRIM28) **(Fig. 5, 6, Data S1-6)**. Our UL78 proximity-dependent labelling experiment also identified several viral transcriptional regulators including the activators: UL26, UL35, UL49, UL69, UL72, UL82, UL95, UL112, and UL122; and repressors: UL34, UL84, UL138. HCMV leverages components of the DNA damage response, particularly the ataxia-telangiectasia mutated (ATM) and ataxia telangiectasia and Rad3-related (ATR) kinase pathways, to facilitate chromatin remodeling and promote the transcriptional activation of the major immediate early promoter (69,70). These kinases phosphorylate downstream effectors such as H2AX and checkpoint kinase 2 (Chk2) to create a permissive environment for viral gene expression (71,72). In parallel, RNA transport and metabolism are tightly regulated during HCMV infection to support the efficient processing and nuclear export of viral transcripts. HCMV lytic infection is associated with altered expression and activity of RNA-binding proteins and splicing factors, such as ribonuclear proteins (RNPs) and serine/arginine-rich (SRs) proteins, which enhance the stability and translational competence of viral mRNAs (73,74). Furthermore, viral proteins like UL69, identified here as a potential UL78 interactor, mimic host mRNA export factors to promote the nuclear export of viral transcripts, bypassing typical cellular restrictions (75–77). Future studies will aim to determine if UL78 can directly or indirectly modify the viral genome to promote transcription during reactivation.

Work presented here, and by others (22), has identified that the bulk of UL78 protein is localized to the plasma membrane and endosomal pools, but nuclear localization has also been suggested in the context of infection (19). Our biochemical approaches to quantify the sub-cellular localization of UL78 using the HiBiT tag fused to the N’ terminus of UL78 determined that between 25-30% of total UL78 (both WT and the DAL mutant) is found at the plasma membrane during infection of human fibroblasts **(Fig. 3B)** with a smaller fraction present in nuclear fractions (**Fig. 7,8**). We also observed a fraction of UL78 associated with the nuclear membrane by confocal microscopy. The capacity to label cellular and viral proteins found within the nucleus as well as the nuclear pore complex suggests that UL78 is oriented such that the C-terminal tail is within the nucleoplasm. There are several ways that GPCRs can traffic to the nucleus including: (1) agonist-dependent or -independent endocytosis from the cell surface and transport to the nucleus mediated by importins, Rabs and sorting nexins (SNX) and (2) agonist-independent translocation from the trans-Golgi network to the nucleus via Rabs and importins (**Fig. 9**). While a specific ligand for UL78 has not been identified to date, a recent structural analysis suggests that the N’ terminus of the protein may in fact cover the ligand tunnel area to either self-activate or prevent access of ligands, which makes it currently challenging to assign ligand-dependent vs. ligand-independent nuclear translocation mechanisms for UL78 (21). Our data showing that the UL78-DAL mutant is also found in the nucleus would imply that ligand-dependent signaling may not be required for nuclear localization. Our UL78 proximity-dependent labeling experiments identified several Rab GTPases and SNX proteins as candidate interaction partners that are likely involved in initial endocytosis and recycling (Rabs 3B, 8A, 21, 35 and SNXs 5, 6, 9 and 18). Intriguingly, Rab11a and SNX11 have been implicated in the translocation of cellular GPCRs from the plasma membrane to the nucleus (57,64,78) and were enriched in our datasets. Alternatively, a fraction of UL78 may be transported to the nucleus directly from the endoplasmic reticulum without trafficking to the cell surface. We identified importin proteins as significantly enriched within our datasets, providing a direct mechanism for the import of UL78 into the host cell nucleus. Two putative classical nuclear localization signals exist within the C’ terminal tail of UL78 at positions 328 – 336 and 360 – 365, suggesting candidate interaction sites for import machinery. It is also possible that UL78 interacts with other cellular proteins that facilitate its nuclear localization. While for now the precise mechanisms of UL78 nuclear translocation are unclear, further host and viral genetic studies could uncover how UL78 gets to the nucleus.

Of the four HCMV-encoded GPCRs, two have previously been investigated in the context of latency and reactivation in CD34^+^ HPCs. UL33 is necessary for virus reactivation through the phosphorylation and activation of the transcription factor CREB (8). We, and others, have shown that US28 expression is necessary for virus latency and reactivation *in vitro* and *in vivo* in humanized mice in a ligand-dependent manner that involves signaling through specific G proteins and downstream effectors (34–36,79). In fact, the multiple roles for US28 at different stages of HPC infection highlight the complexity of GPCR signaling during viral infection of progenitor cells. Herein we provide a role for UL78 in reactivation from latency in CD34^+^ HPCs and indicate that Ga_i_ coupling is important for this phenotype. Our work also uncovers the nuclear localization of a fraction of UL78, although it is still not known whether this process is essential for efficient reactivation from latency. These findings add to the growing appreciation of the role of virally-encoded GPCRs play in latency and reactivation in CD34^+^ HPCs. UL78 awaits the identification of a ligand, either extracellular or intracellular, and a more mechanistic understanding of its function(s) in different cellular compartments which will lead to a greater understanding into the role played by UL78 in promoting virus reactivation in hematopoietic cells.

## Materials & Methods

### Plasmids

Plasmids were generated utilizing traditional cloning methodology as previously described using primers listed in **Table S1** (9). Briefly, HA-UL78-NP was generated by cloning the Natural Peptide (NP) tag (GVTGWRLCERILA) in-frame with the C-term of UL78. HB-UL78 was generated by cloning the HiBiT (HB) tag (VSGWRLFKKIS) in-frame with the N-term of UL78. Fragments were PCR amplified and cloned into the pcDNA3.1-vector. Mutations in the DRL motif of UL78 were generated by site-directed mutagenesis, substituting alanine for the indicated amino acid using the Q5 Mutagenesis Kit (NEB) following the manufacturer’s recommended procedure. All constructs were confirmed by sequencing and transformed into TOP10 Escherichia coli cells (Invitrogen). Large-BiT tagged Gα subunits were kindly provided by Julien Hanson (Addgene plasmid ID: 134359, 134360, 134364, and 134363) (47).

### Cells and Virus

Normal human dermal fibroblasts (NHDFs) were obtained from ATCC (PCS-2021-010) and human embryonic kidney (HEK) −293 cells were obtained from Microbix. NHDF and HEK-293 cells were maintained in Dulbecco’s modified eagle’s medium (DMEM) supplemented with 10% fetal bovine serum (FBS), streptomycin, penicillin, and glutamine at 37° C and 5% CO_2_. M2-10B4 and S1/S1 stromal cells were obtained from Stem Cell Technologies and cultured as previously described (80). WA01 human embryonic stem cells (hESCs) were obtained from the WiCell Research Institute – National Stem Cell Bank and were cultured as previously described (34,40). The HCMV strain TB40/E-GFP, which constitutively expresses green fluorescent protein under the SV40 promoter (81), was amplified in NHDFs as previously described (53,82). Recombinant viruses were generated using a two-step recombineering procedure utilizing the HCMV TB40/E-GFP bacterial artificial chromosome (BAC). Viral constructs were confirmed by next generation sequencing prior to plaque purification and clonal expansion. Viral titers were determined via plaque assay on NHDF cells and aliquots stored at −80° C. For viral growth analyses, single-step growth curves were carried out at a multiplicity of infection (MOI) of 3.0 PFU/mL, and multi-step growth curves were carried out at a MOI of 0.01 PFU/mL. Supernatant and cell-associated virus were harvested at multiple time points post-infection and titered via limiting dilution plaque assay on NHDF cells.

### Immunoblot

Blotting procedures were carried out as previously described (9,34). Briefly, cell lysates were harvested using either RIPA Lysis Buffer (Santa-Cruz Biotechnology) supplemented with HALT protease inhibitor (Thermo-Fisher Scientific) or utilizing the NE-PER extraction kit (ThermoFisher Scientific). Proteins were separated on a 4-12% SDS-PAGE gel and transferred onto PVDF membranes. Immunoblots were performed using antibodies directed against β-Actin (sc-47778, Santa-Cruz Biotechnology), HCMV IE1/IE2 (MAB8131, Millipore-Sigma), HA (sc-7392, Santa-Cruz Biotechnology), HCMV gB (sc-69742, Santa-Cruz Biotechnology), Streptavidin-HRP (21130, Thermo Scientific), NUP98 (C39A3, Cell Signaling Technology), ERp57 (CL2444, Thermo Scientific), GAPDH (sc-47724, Santa-Cruz Biotechnology), and, if required, the appropriate HRP-conjugated secondary antibody (sc-525409, Santa-Cruz Biotechnology). HiBiT-tagged proteins were visualized using the Nano-Glo HiBiT Blotting System (Promega).

### Proximity-Dependent Labeling Experiments

Proximity-dependent labeling experiments were conducted as previously described (34). Briefly, monolayers of NHDF cells or CD34^+^ HPCs were either mock infected, or infected HCMV TB40/E-GFP-UL78-TurboID or TB40/E-GFP at an MOI of 2. For experiments utilizing NHDFs, at three days post infection, cells were incubated for 6 hours in complete media supplemented with 50μg/mL biotin. Cells were either lysed in RIPA buffer (50 mM Tris pH 8, 150 mM NaCl, 1% triton x-100, 0.1% SDS) and 1x Halt protease inhibitor cocktail (ThermoFisher) and centrifuged at 10,000 relative centrifugal force at 4°C or were processed utilizing the NE-PER extraction kit (ThermoFisher Scientific). For experiments utilizing CD34^+^ HPCs, cells were cultured in LTBMC as previously described (80). At 14 days post-infection, HPCs were placed into RPMI-1640 medium containing 20% FBS, 2 mM L-glutamine, 100 U/mL penicillin, 100 μg/mL streptomycin, 15 ng/mL granulocyte-colony stimulating factor (G-CSF), 15 ng/mL granulocyte-macrophage colony-stimulating factor (GM-CSF), and 50 µg/mL of biotin and overlaid onto confluent monolayers of NHDFs for 16 hours prior to cell lysis. Resultant lysates were incubated with 250μL Pierce NeutrAvidin Agarose beads (ThermoFisher) overnight at 4°C while rotating. Beads were collected and washed several times using urea wash buffer (PBS pH 7.4, 4 M urea), wash buffer 2 (PBS pH 7.4, 1% triton x-100), 50 mM ammonium bicarbonate, and 6 M urea. Proteins were reduced and alkylated using 0.5 M tris(2-carboxyethyl)phosphine and 0.5 M iodoacetamide prior to tryptic digestion. Digestion was halted by adding 20 µL formic acid to each sample and samples were stored at −80° C until LC-MS/MS.

### LC-MS/MS and Data Analysis

LC-MS/MS was performed as previously described by the Fred Hutchinson Proteomics Core (Seattle, WA) (34). Briefly, samples were desalted using ZipTip C18 (Millipore, Billerica, MA) and eluted with 70% acetonitrile/0.1% TFA (Trifluoracetic acid; Sigma) and the desalted material dried in a speed vac. Desalted samples were brought up in 2% acetonitrile in 0.1% formic acid (12 μL) and 10 μL of sample analyzed by LC/ESI MS/MS with a ThermoScientific Easy-nLC II nano HPLC system (Thermo Scientific, Waltham, MA) coupled to a tribrid Orbitrap Fusion mass spectrometer (Thermo Scientific, Waltham, MA). Peptide separations were performed on a reversed-phase column (75 μm × 400 mm) packed with Magic C18AQ (5-μm 100Å resin; Michrom Bioresources, Bruker, Billerica, MA) directly mounted on the electrospray ion source. A 90-minute gradient from 7% to 28% acetonitrile in 0.1% formic acid at a flow rate of 300 nL/minute was used for chromatographic separations. The heated capillary temperature was set to 300°C and a static spray voltage of 2100 V was applied to the electrospray tip. The Orbitrap Fusion instrument was operated in the data-dependent mode, switching automatically between MS survey scans in the Orbitrap (AGC target value 500,000, resolution 120,000, and maximum injection time 50 milliseconds) with MS/MS spectra acquisition in the linear ion trap using quadrupole isolation. A 2 second cycle time was selected between master full scans in the Fourier-transform (FT) and the ions selected for fragmentation in the HCD cell by higher-energy collisional dissociation with a normalized collision energy of 27%. Selected ions were dynamically excluded for 30 seconds and exclusion mass by mass width +/− 10 ppm.

Data analysis was performed using Proteome Discoverer 2.2 (Thermo Scientific, San Jose, CA) searching against the UniProt Human (proteome ID: UP000005640) and HCMV TB40/E (proteome ID, UP000143167) proteomes. Trypsin was set as the enzyme with maximum missed cleavages set to 2. The precursor ion tolerance was set to 10 ppm and the fragment ion tolerance was set to 0.6 Da. Variable modifications included oxidation on methionine, carbamidomethyl on cysteine, and acetylation on protein N-terminus. Normalized LFQ intensities were inputted into Perseus (83) where proteins with greater than 70% missing values were removed. Remaining missing values were imputed from the normal distribution. Protein abundances were Log_2_ transformed, and a student’s t test corrected for multiple comparisons was performed. Proteins were considered candidate interaction partners of HCMV UL78 if the CRAPome frequency was <30%, FDR-corrected q value <0.05, and the Log_2_ fold change was > 1.5.

### Live Cell G-Protein Coupling Assay

Receptor G-protein coupling was assessed as previously described (9,47) using the Nano-Glo Live Cell Assay system (Promega). Briefly, HEK-293 cells were seeded into treated black 96-well plates at a density of 3.0×10^4^ cells per well. The following day, cells were co-transfected in triplicate with a 1:1 ratio of the indicated GPCR constructs or the empty pcDNA3.1-vector, and Large BiT linked Gα subunit using Fugene4K (Promega) following the manufacturer’s recommended procedure. At 18 hours post-transfection, the growth medium was replaced with Opti-MEM media (Thermo Fisher Scientific). At 6 hours post media replacement, 25 μL of reconstituted Nano-Glo Live Cell assay reagent was added to each well, and plates were briefly incubated with agitation. Luminescence, indicative of G protein coupling, was measured using a Promega GloMax Navigator luminometer. Assay results were transferred to a Microsoft Excel spreadsheet, backgrounds subtracted and plotted in GraphPad Prism 10.0 software.

### HiBiT Split-Luciferase Assay

For surface vs total expression assays, NHDFs were seeded into cell culture-treated black 96-well plates at a density of 1.5×10^4^ cells per well. The following day, triplicate wells were infected with the indicated HiBiT-tagged recombinant viruses at a MOI of 1.0. At 72 hpi, surface vs. total HiBiT expression was evaluated using the Nano-Glo HiBiT Extracellular and Lytic Detection kits (Promega) following the manufacturer’s recommended procedure. For nuclear localization assays, NHDFs were seeded into cell culture-treated 6-well plates at a density of 3.0×10^5^ cells per well. The following day, replicate wells were infected with the indicated HiBiT-tagged recombinant viruses at a MOI of 1.0. At 72 hpi, cytoplasmic and nuclear extracts were harvested utilizing the NE-PER extraction kit (ThermoFisher Scientific). Lysates were plated in triplicate into black 96-well plates and HiBiT expression was evaluated using the Nano-Glo HiBiT Lytic Detection kits (Promega) following the manufacturer’s recommended procedure. Luminescence was measured using a Promega GloMax Navigator luminometer. Assay results were transferred to a Microsoft Excel spreadsheet, background subtracted, normalized to the HiBiT control protein, and % surface expression was determined by using the ratio of extracellular vs lytic luminescence. Results were analyzed using GraphPad Prism 10.0 software.

### Microscopy

NHDFs were grown on 13mm glass coverslips and infected at an MOI of 0.5 with WT-HCMV, HB-UL78, or HB-UL78-DAL. At 72 hpi, coverslips were washed with PBS and fixed with 4% paraformaldehyde in PBS. Cells were permeablized with 0.25% Triton, blocked with normal goat serum, and stained with the indicated antibodies. Coverslips were then washed with PBS and incubated with the appropriate fluorophore-conjugated secondary antibodies. Fluorescence was visualized using a LEICA Stellaris 8 microscope using a 63x objective with an NA of 1.4. The fluorophores were excited using 405nm and White Light Lasers. The signals were captured using Leica Stellaris 8 and the Leica Application Suite Software. Images were exported as .tiff files and analyzed using ImageJ software.

### HCMV Latency and Reactivation Assay

hESC-derived CD34^+^ HPCs were differentiated from WA01 human embryonic stem cells using the commercial STEMdiff Heme feeder-free hematopoietic differentiation kit (Stem Cell Technologies) as previously described (40,52,82). HPCs were cultured in IMDM with 10% BIT serum replacement, stem cell cytokines (stem cell factor, FLT3L, IL-3, and IL-6 [PeproTech]), and penicillin/streptomycin as previously described (11,53,82). CD34^+^ HPCs were infected with the indicated viruses at a MOI of 2 for 48 hours prior to isolation by fluorescence-activated cell sorting (FACS) using a FACSAria (BD FACS Aria equipped with 488, 633, and 405 nm lasers, running FACS DIVA software) in order to obtain a pure population of viable, GFP^+^, CD34^+^, HPCs as previously described (80). Infected cells were co-cultured in transwells above monolayers of irradiated M2-10B4 and S1/S1 stromal cells. At 14 days post-infection, HPCs were serially diluted in RPMI-1640 medium containing 20% FBS, 2 mM L-glutamine, 100 U/mL penicillin, 100 μg/mL streptomycin, 15 ng/mL granulocyte-colony stimulating factor (G-CSF), and 15 ng/mL granulocyte-macrophage colony-stimulating factor (GM-CSF) and overlaid onto confluent monolayers of NHDFs cultured in 96-well plates in an extreme limiting dilution assay. To quantify the levels of pre-reactivation infectious virus, a fraction of the HPC cultures were mechanically disrupted and then used in the ELDA. Cell cultures were microscopically visualized for the presence of GFP^+^ weekly, for up to 3 weeks, and the frequency of infectious center production was calculated by ELDA software (41).

### Viral DNA Quantification

Primers and probes recognizing HCMV UL141 were used to quantify viral genomes by quantitative real-time PCR (11). Briefly, total DNA was extracted using Trizol (ThermoFisher) according to the manufacturer’s recommendations. Dilutions of purified HCMV BAC DNA were used to create a standard curve. Total DNA was added to each reaction well of TaqMan FastAdvance PCR master mix (Applied Biosystems) and samples were analyzed in triplicate on a StepOnePlus TaqMan PCR machine (Applied Biosystems) with an initial activation at 50°C for 2 min and 95°C for 20 s, followed by 40 cycles of 1 s at 95°C and 20 s at 60°C. TaqMan results were analyzed using ABI StepOne software and graphed using GraphPad Prism 10.0 software.

## Acknowledgments

This work was supported by grant P01 AI127335 from the National Institute of Allergy and Infectious Diseases (NIAID), National Institutes of Health (NIH), awarded to P.C., M.H.H., and D.N.S. S.M. is supported by National Institutes of Health (NIH) T32AI170496-01A1. M.H.H. and P.C. are supported by National Institutes of Health (NIH) R37 AI21640. This research utilized the Integrated Pathology Core at the Oregon National Primate Research Center (ONPRC) which is supported by NIH Award P51 OD 011092. The funder had no role in study design, data collection and analysis, decision to publish, or preparation of the manuscript.

## Author Contributions

S.M., N.L.D., D.M., P.C., D.N.S. and M.H.H. designed research; S.M., N.L.D., M.D., R.L.T., L.J.P., A.T.M., J.M., L.S., L.H., T.B., G.S., C.N.K., D.M., P.C., D.N.S., and M.H.H. performed research; S.M., N.L.D., M.H.H., P.C., and D.N.S. analyzed data; and S.M., N.L.D., M.D., D.N.S., and M.H.H. wrote the paper with input from all authors. The authors declare no competing interest.

